# Diversity and functional analysis of rumen and fecal microbial communities associated with dietary changes in crossbreed dairy cattle

**DOI:** 10.1101/2022.08.29.505658

**Authors:** Felix M. Kibegwa, Rawlynce C. Bett, Charles K. Gachuiri, Eunice Machuka, Francesca Stomeo, Fidalis D. Mujibi

**Affiliations:** Department of Animal Production, Faculty of Veterinary Medicine, University of Nairobi, Nairobi, Kenya; Biosciences eastern and central Africa - International Livestock Research Institute (BecA-ILRI) Hub, Nairobi, Kenya; Genomics Division, USOMI Limited, Nairobi, Kenya

**Keywords:** Rumen metagenome, fecal metagenome, dairy cattle, concentrates, diet

## Abstract

The objective of this study was to investigate the effect of varying roughage and concentrate proportions, in diet of crossbreed dairy cattle, on the composition and associated functional genes of rumen and fecal microbiota. We also explored fecal samples as a proxy for rumen liquor samples. Six crossbred dairy cattle were reared on three diets with an increasing concentrate and reducing roughage amount in three consecutive 10-day periods. After each period, individual rumen liquor and fecal samples were collected and analyzed through shotgun metagenomic sequencing. Average relative abundance of identified Operational Taxonomic Units (OTU) and microbial functional roles from all animals were compared between diets and sample types (fecal and rumen liquor). Results indicated that dietary modifications significantly affected several rumen and fecal microbial OTUs. In the rumen, an increase in dietary concentrate resulted in an upsurge in the abundance of Proteobacteria, while reducing the proportions of Bacteroidetes and Firmicutes. Conversely, changes in microbial composition in fecal samples were not consistent with dietary modification patterns. Microbial functional pathway classification identified that carbohydrate metabolism and protein metabolism pathways dominated microbial roles. Assessment of dietary effects on the predicted functional roles of these microbiota revealed that a high amount of dietary concentrate resulted in an increase in central carbohydrate metabolism and a corresponding reduction in protein synthesis. Moreover, we identified several microbial stress-related responses linked to dietary changes. *Bacteroides* and *Clostridium* genera were the principal hosts of these microbial functions. Therefore, the roughage to concentrate proportion has more influence on the microbial composition and microbial functional genes in rumen samples than fecal samples. As such, we did not establish a significant relationship between the rumen and fecal metagenome profiles, and the rumen and fecal microbiota from one animal did not correlate more than those from different animals.

## Introduction

Demand for animal source foods is rapidly steadily rising: for example, milk demand in low-income countries is expected to increase by 136% by 2030 compared to 2000 (1). This increase in demand has been ascribed mostly to population expansion, increased urbanization, and income growth (2,3). To meet this huge demand, developing countries have to significantly increase livestock production, (3– 5), through simultaneous interventions in animal feeds, genetics, and health (6).

In East Africa, many interventions to improve milk production have had minimal gains because farmers depend mainly on rain-fed forage, pastures/cultivated fodder and crop by-products. Additionally, the high cost of conventional feed resources, such as cereals and grain legumes, the food-feed competition between man and livestock, and the absence of suitable technology to optimize the use of these feeds in animal husbandry prohibit their wide-scale use (7). On the other hand, despite continuous improvement of the cattle genotypes through cross-breeding and upgrading, the proportion of improvement in milk production attributable to this genetic improvement remains relatively low (8). There is therefore a need for a paradigm shift that incorporates use of new technologies or customization of existing technologies to improve milk production while using the available feed resources and cattle genotypes. Previous studies have found that animal’s diet (quantity and quality) is closely related to animal production and the composition of rumen microbes (9,10). This is because the rumen microbiota symbiotically degrades forages into nutrients, such as volatile fatty acids and microbial proteins, which are utilized by the host animal (11,12). However, few studies have been conducted to evaluate the microbial composition in crossbred cattle reared by dairy farmers in the East African tropics and subjected to constant fluctuations in diet composition.

To facilitate rumen microbial composition evaluation, the standard sample collection method is rumen cannulation of cattle (13). However, only a small number of ruminally cannulated cattle are accessible to any investigator in a single study, thus restricting the number of cattle that can be used to assess the ruminal microbiome, if only rumen cannulated cows are used. A more conventionally accepted method of collecting rumen contents from non-cannulated cattle is using stomach tubing. A study by Paz *et al*. (13) showed that samples collected from the same rumen cannulated animals and stomach tubing did not result in a significant difference in the composition of ruminal microbiome. However, this approach cannot be used to repeatedly collect rumen samples from the same cattle within a short period because the cattle are stressed during the sampling (14). Additionally, the sampling technique requires the use of expensive equipment and well-trained personnel. It is therefore important to explore other potential non-invasive alternative samples/sampling techniques that can act as proxies for assessing the composition of the rumen microbial community.

For many years, the structure of rumen microbial community has been evaluated by the culturing of samples using selective media (15–17). However, this method is time-consuming and heavily biased by the selected isolation media and methods employed, thus providing an incomplete depiction of the microbial community structure (10,18). The advent of modern genomics and molecular microbial techniques, such as next generation sequencing (NGS), has facilitated in-depth sequencing and analysis of environmental specimens, including the rumen microbiota, at a higher and more precise level than the previous methods (19,20). NGS technology is different from the Sanger sequencing method because of its massively parallel sequencing approach, high throughput, and cost reduction. This technology allows for longer read lengths, faster turn-around time, more reads per unit cost, and reduced error rates (21). These advantages of NGS has led to its substantial use in metagenomic studies (22–26).

Accordingly, we hypothesized that given the advancements in NGS technology and the sequencing depth of shotgun sequencing, it would be possible to detect even the slightest changes in microbial composition and microbial functional genes across different sections of the cattle gastrointestinal (GI) tract. Therefore, the objective of this study was to determine the effect of varying roughage and concentrate proportions, in diet of crossbreed dairy cattle, on the composition and associated functional genes of rumen and fecal microbiota. Additionally, rumen liquor and fecal metagenome profiles were contrasted to determine if the fecal metagenome was predictive of rumen metagenome.

## Materials and Methods

### Animals and diets

This study was approved and performed in accordance with the guidelines of University of Nairobi’s Faculty of Veterinary Medicine Animal care and use committee (ACUC). Animals were handled by experienced animal health professionals to minimize discomfort and injury.

Experimental animals were chosen from the dairy herd of the Faculty of Veterinary Medicine Farm at the University of Nairobi (latitude 1° 14′33.4″S, 36° 42′36.3″E). The experimental design was discussed in our previous study (27). This research used six crossbred lactating cows (300 ± 50 kg body weight; 174 ± 15 days in milk) in first-parity. Before the experiments (acclimatization period), all the animals were fed *ad libitum* mixed diet of *Pennisetum clandestinum* (Kikuyu grass) and *Chloris gayana* (Rhodes grass) hay, supplemented with dairy meal (a commercial concentrate), at 0.01/kg body weight. The feeds were divided into two and offered during morning and evening milking. The animals were then fed on three distinct diets in three successive 10-day periods after this 10-day acclimatization period. The experimental diets contained an increased amount of concentrate and a reduced amount of roughage. The roughage ratios to concentrate in the diets were 90:10 (Diet 1), 75:25 (Diet 2), and 60:40 (Diet 3). The three diets were developed using the NRC-Nutrient Requirements of Dairy Cattle Software v 1.9 (28) to meet the energy demands of dairy cows producing 20 kg of milk/day with 4.0 % milk fat and 3.5 % true protein. For the roughage, the dietary components were *Chloris gayana* (Rhodes grass), hay, *Pennisetum purpureum* (Napier grass), *Pennisetum clandestinum* (Kikuyu grass), and dry *Zea mays* (maize) Stover. These were mixed with dairy meal and urea in different proportions to make total mixed rations of the three diets, as shown in Table A in S1 Text, and offered to the animals at 8 am and 6 pm. The chemical composition of the dietary ingredients was evaluated according to the Association of Official Analytic Chemist (AOAC) methods (29), while a previously described method (30) was used to assess dietary fiber (Table B in S1 Text). The cattle were housed in individual stalls and had free access to fresh water and mineral lick supplements throughout the 30 days of the experiment.

### Sample collection and DNA extraction

A total of 36 (18 fecal and 18 rumen liquor) samples were collected in this study. The samples were obtained from individual animals on the final day of each experimental diet, i.e., day 10 (for diet D1), day 20 (for diet D2), and day 30 (for diet D3). Roughly, 200 g of fecal samples were obtained by rectal grab, and a sub-sample was placed into a sterile 50 mL falcon tube. As previously described, rumen liquor samples were collected by a suction pump and a flexible stomach tube (31). Approximately 250 mL of total (i.e., solid and liquid) rumen contents were collected, and 50 mL of the rumen liquor was placed in sterile 50 mL falcon tubes after discarding the first 200 mL of sample to reduce saliva contamination. The samples were transported to the Biosciences eastern and central Africa-International Livestock Research Institute, (BecA ILRI) Hub laboratory in Nairobi, and stored at-20°C awaiting further analysis. All samples were kept in a vaccination cooler box with dry ice during the transportation from the field to the laboratory.

Before DNA extraction, samples were thawed at room temperature (∼22°C) and thoroughly mixed with vortexing at maximum speed for 30 seconds. Total DNA was extracted from a representative subsample of fecal and rumen liquor samples using a QIAamp^®^ DNA Stool Mini Kit (Qiagen, Valencia, CA, USA), following the manufacturer’s guidelines with a few changes. The modifications were made to increase the DNA yield while reducing the amount of RNA obtained. The modifications included (i) doubling the recommended amount of the sample and (ii) adding 2 µl of RNAse A to the sample and proteinase K mixture. After elution, the DNA was visualized with UV light on 1 % agarose gel electrophoresis. The DNA quantity and quality were assessed by using the NanoDrop spectrophotometer (ThermoScientific, USA) and Qubit 2.0 Fluorometer (Thermo Fisher Scientific, USA), following the manufacturer’s recommendations.

### Preparation of DNA Library and Illumina MiSeq sequencing

The Nextera DNA Preparation Kit and the Nextera Index Kit (Illumina, San Diego, CA, USA) were used to prepare the DNA Libraries following the manufacturer’s instructions with a few modifications. About 50 ng of total genomic DNA per sample underwent simultaneous tagmentation and the addition of adapter sequences at 55 °C for 10 min. The resulting tagmented mixture was purified using the Zymo DNA Kit (Zymo Research Corporation, Irvine, CA, USA). The Zymo IIC spin columns were used, and all the centrifugation steps were performed at 10,000 ×g, not following the manufacturers’ recommendation that suggested using a Zymo-Spin I-96 Plate centrifuged at 1300×g. A limited-cycle (5 cycles) polymerase chain reaction (PCR) was then conducted to amplify the insert DNA using a unique combination of barcode primers. This PCR reaction also added index sequences on both ends of the DNA. Finally, PCR products were cleaned up, and short library fragments, including indexes, were removed using AMPure XP beads (A63881, Beckman Coulter, Brea, CA, USA). The final library concentration was measured using the Qubit dsDNA HS Assay Kit (Thermo Fischer Scientific, USA) and the average library size was estimated using the Bioanalyzer TapeStation 2200 (Agilent Technologies, Santa Clara, USA). The libraries were diluted to 4 nM stocks, pooled in equimolar ratios, spiked with 1 % PhiX, and paired-end (200 cycles) sequenced on the MiSeq^®^ (Illumina, USA) platform at the BecA-ILRI Hub, Nairobi, Kenya.

### Analysis and processing of sequence reads

The quality of all paired-end raw fastq sequencing reads were evaluated using FastQC (version 0.11.5) (http://www.bioinformatics.babraham.ac.uk/projects/fastqc/). Before further sequence analysis, filters were used to remove reads with poor quality from all the samples. SolexaQA v3.1.5 (32) was used to calculate sequence quality statistics and perform quality filtering of the raw reads. Reads were then trimmed at a threshold of Q20 using DynamicTrim in SolexaQA++. Filtered reads were re-assessed for quality using FastQC, and any samples that still had a poor quality were further processed by truncating at any site having an average quality score < 20 using the FASTX-trimmer in the FASTX-toolkit v0.0.14 (http://hannonlab.cshl.edu/fastx_toolkit/). The cleaned fastq sequences were uploaded onto the publicly available Meta Genome Rapid Annotation using the Subsystem Technology server (MG RAST, v3.3) (33). The taxonomic domain groups were allotted using MG RAST against the M5NR database. The data were also analyzed using the SEED Subsystems platform in MG RAST to identify putative protein-encoding sequences. SEED subsystems platform annotates the sequences in a set of end-to-end homologous proteins that share a common function (FIGfams), based on the FIGfam database, then maps these protein families against the SEED subsystems to infer metabolic pathways in what is called subsystem reconstruction. This functional classification in SEED hierarchical classification had a percentage identity cut-off of 60 %, an expected value (E) cutoff of 10^−5^, and a minimum alignment cut-off of 15 bp (34).

### Statistical analysis

The operational taxonomic unit (OTU) counts were normalized by relative abundance and log transformation, [log(*x* + 1)], before quantitative characterization using the Paleontological STatistics software package (PAST v3.13), (35). Alpha diversity analysis was conducted in PAST v3.13 to evaluate the taxonomic richness and diversity using Chao1 minimal richness index (Chao and Shen, 2003), inverse Simpson diversity index (37,38), and Shannon diversity index (39). One-way analysis of variance (ANOVA) was also performed to assess individual OTU differences per diet within each sample type using Genstat version 14 (40), and correction for multiple testing was performed using the Bonferroni adjustment. To group sample types according to their characteristics, cluster analysis of the metagenomes was performed using principal component analysis (PCA). The significance of the sample type was calculated using one-way PERMANOVA on 9,999 permutations (p < 0.01). Linear discriminant analysis Effect Size (LEfSe) (41), was used in the specific identification of OTUs that differed between the two sample types (fecal and rumen liquor). This approach uses a non-parametric factorial Kruskal-Wallis sum-rank test and a linear discriminate analysis to detect both statistically significant and biological germane features. The core OTUs relative abundances were used as an input for LEfSe. The LefSe analysis also used to identify specific OTUs that differed between sample types. The LEfSe analysis conditions were set as follows: i) alpha value for the factorial Kruskal-Wallis test among classes at < 0.05; ii) alpha value for the pairwise Wilcoxon test among subclasses at < 0.05; iii) the threshold on the logarithmic LDA score for discriminative features at < 2.0; and iv) multi-class analysis was set as all-against-all. Additionally, two-group analysis was performed, applying a Fisher’s exact *t*-test with a 95 % confidence interval to assess the differences in the abundance of microbes between fecal and rumen liquor samples. Pearson correlation analysis was performed with Genstat (version 14) to evaluate the relationship between fecal and rumen liquor samples within and between animals. Next, a *t*-test was performed on the correlation values to weigh if the correlation between the rumen and fecal profiles of a cow was more significant than the correlation between samples from different cows. Finally, to estimate the differences in functional pathways, SEED Level 1 and 2 functional classification, one-way ANOVA to assess dietary effects, and a t-test to evaluate sample type effects were conducted using Genstat.

## Results

### Sequence Assessment

Metagenomic sequencing of the samples resulted in a total of 9.6 and 11.9 million raw reads for fecal and rumen liquor samples, respectively. After quality control, the total number of reads was reduced by 27.63 % for fecal samples and 26.39 % for rumen liquor samples. Upon MG-RAST annotation, less than 1 % of reads per diet for both fecal and rumen liquor sample types were classified as rRNA based on hits against 16S rRNA gene sequence databases. About 73.56–81.22 % and 74.54–78.32 % of the reads from fecal and rumen liquor samples, respectively, were classified into various functional subsystems (Table C in S1 Text). The classification of individual OTUs identified 55 phyla, 131 classes, 253 orders, 443 families, and 901 genera in four domains. Given the large number of classified groups and to facilitate the interpretation of results, our investigation focused on the most abundant groups within each domain. The bacteria domain was dominant irrespective of sample type and diet. Together, the domains of archaea, Eukaryota, and virus comprised less than 2 % of all the sequences found in both fecal and rumen metagenomes (Fig 1). Moreover, roughly 1 % of the reads could not be categorized into any recognized OTU.

**Fig 1.**
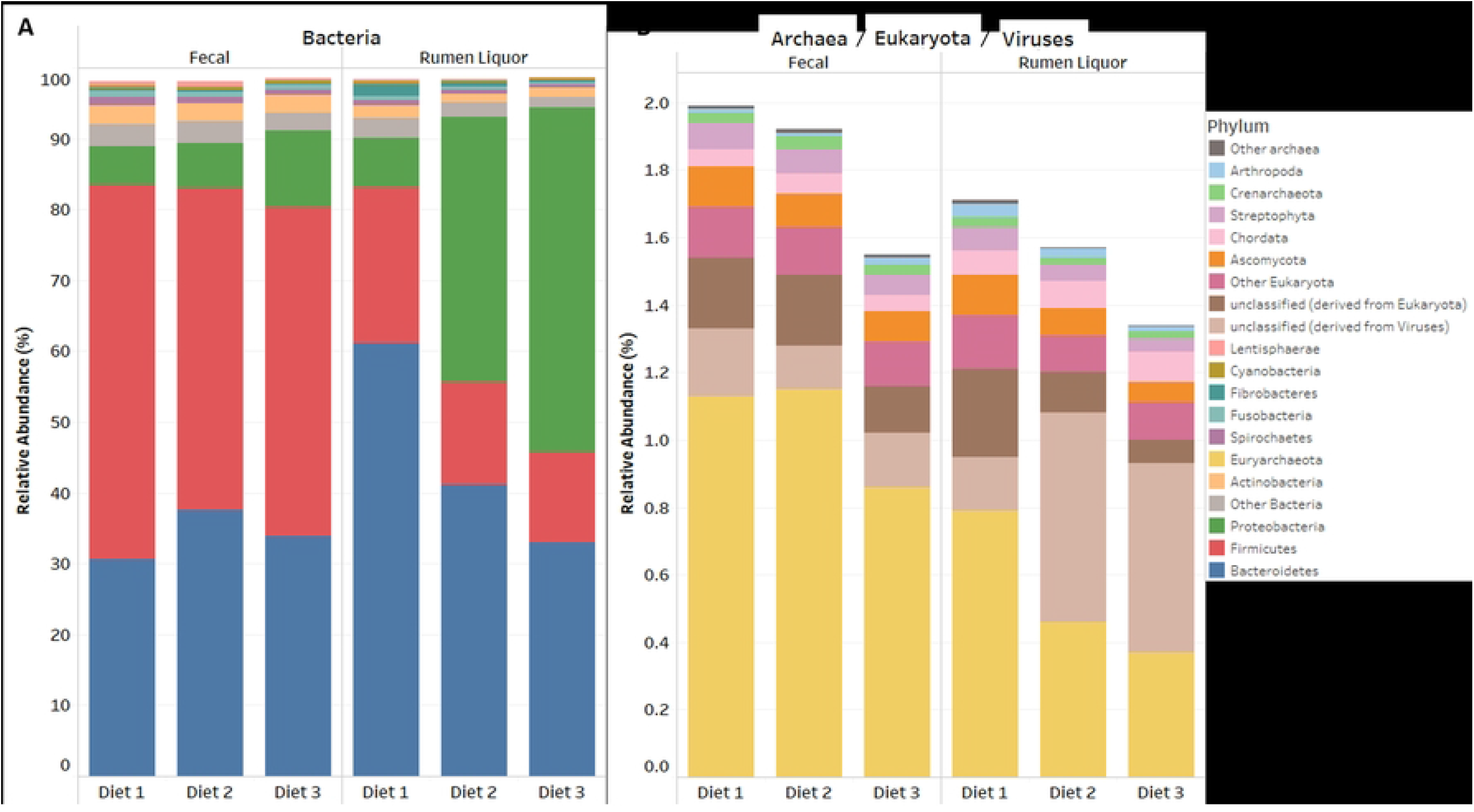
Circular bar plot of relative abundance of all phyla in the three diets. Abundances outside the circle are from rumen liquor samples, while abundances inside the circle are from fecal samples. **A** is for archaea, **B** is for bacteria, **E** is for eukaryota, and **V** is for virus domains. The three most abundant phyla are highlighted by the red trapezoid.

### Effect of diet on the diversity and relative abundance of fecal and rumen microbiota

Table 1 shows the estimators of diversity and evenness within each diet in fecal and rumen liquor sample types. ANOVA revealed no significant differences, in all diversity and evenness indices, between diets for both rumen liquor and fecal samples. However, the principal component analysis (PCA) undertaken on the entire dataset of microbial counts of fecal and rumen liquor samples indicated clustering of samples based on the diet in rumen liquor samples as opposed to fecal samples (Fig 2). Similarly, when the relative abundance of each taxon was compared, a significant effect of diet on the composition of the microbial community in several phyla and genera was observed (Table 2). Within fecal samples, Firmicutes (p = 0.02), Proteobacteria (p = 0.01), Actinobacteria (p = 0.01), Cyanobacteria (p = 0.01), Fibrobacteres (p = 0.01), Streptophyta (p = 0.04), other archaea (p = 0.04), other Eukaryota (p = 0.01), and unclassified derivatives of viruses (p = 0.01) were significantly different among the diets. Of these significantly different phyla, the phyla Firmicutes, Actinobacteria, Fibrobacteria, Streptophyta, Other Eukaryota, and unclassified derivatives of viruses had the highest abundance in Diet 1, while Proteobacteria had the highest abundance in Diet 3, compared with the other diets. The rest of the phyla had the highest abundance with Diet 2. At the genus level, the abundance of six genera, *Methanococcus, Methanosarcina, Clostridium, Eubacterium, Ethanoligenens*, and *Ruminococcus*, was significantly different with diet changes. All these genera, except *Methanosarcina*, had the highest abundance with Diet 1 and lowest abundance with Diet 3 (Table 2).

**Table 1:** Estimators of diversity within each diet for fecal and ruminal fluid samples (Mean ± Standard Error).

|  | Fecal<sup>1</sup> |  |  | Rumen Liquor<sup>1</sup> |  |  | P value |  |  |
| --- | --- | --- | --- | --- | --- | --- | --- | --- | --- |
|  | Diet 1 | Diet 2 | Diet 3 | Diet 1 | Diet 2 | Diet 3 | S | D | SXD |
| **Chao-1** | 1713 ± 69 | 1675 ± 16 | 1729 ± 16 | 1833 ± 9 | 1786 ± 18 | 1756 ± 36 | * | NS | NS |
| **Simpson (1-D)** | 0.993 ± 0 | 0.993 ± 0 | 0.993 ± 0 | 0.954 ± 0.037 | 0.963 ± 0.002 | 0.974 ± 0.002 | ** | NS | NS |
| **Shannon (H)** | 5.78 ± 0.026 | 5.829 ± 0.012 | 5.819 ± 0.028 | 4.876 ± 0.349 | 5.22 ± 0.036 | 5.213 ± 0.069 | *** | NS | NS |
| **Evenness (e<sup>H</sup>/S)** | 0.191 ± 0.009 | 0.203 ± 0.003 | 0.195 ± 0.005 | 0.101 ± 0.017 | 0.104 ± 0.004 | 0.105 ± 0.006 | *** | NS | NS |
| **Equitability (J)** | 0.777 ± 0.005 | 0.785 ± 0.002 | 0.781 ± 0.003 | 0.622 ± 0.046 | 0.697 ± 0.005 | 0.698 ± 0.008 | *** | NS | NS |

*S = Sample type, D = Diet, SXD Sample type by Diet interaction* *NS = Not Significant, \* = Significant at 0.05, \*\* = Significant at 0.01, \*\*\* = Significant at 0.001* <sup>1</sup>Means are based on 6 cows.

**Table 2:** Mean sample (%) of microbial phyla and genus by sample type and diet.

| Sample Type |  | Fecal<sup>1</sup> |  |  |  | Rumen Liquor<sup>1</sup> |  |  |  | Sample Type P value |
| --- | --- | --- | --- | --- | --- | --- | --- | --- | --- | --- |
| Domain | Phylum/Genus | Diet 1 | Diet 2 | Diet 3 | P value | Diet 1 | Diet 2 | Diet 3 | P value |  |
| Archaea | **Crenarchaeota** | 0.03 | 0.04 | 0.03 | 0.08 | 0.03 | 0.02 | 0.02 | 0.01 | NS |
|  | **Euryarchaeota** | 1.13 | 1.15 | 0.86 | 0.06 | 0.79 | 0.46 | 0.37 | <0.01 | * |
|  | *Methanobrevibacter* | 0.31 | 0.19 | 0.13 | <0.01 | 0.16 | 0.07 | 0.06 | <0.01 | NS |
|  | *Methanosarcina* | 0.16 | 0.18 | 0.13 | 0.02 | 0.14 | 0.09 | 0.07 | <0.01 | * |
|  | **Other archaea** | 0.01 | 0.01 | 0.01 | 0.04 | 0.01 | 0 | 0 | 0.02 | NS |
| Bacteria | **Firmicutes** | 52.79 | 45.35 | 46.4 | 0.02 | 22.02 | 14.53 | 12.78 | 0.01 | ** |
|  | *Clostridium* | 16.07 | 13.91 | 13.05 | 0.04 | 5.14 | 3.41 | 2.9 | 0.01 | * |
|  | *Eubacterium* | 4.28 | 3.64 | 3.27 | 0.01 | 2.02 | 1.25 | 1.09 | <0.01 | NS |
|  | *Ethanoligenens* | 1.37 | 1.15 | 1.08 | 0.04 | 0.24 | 0.15 | 0.12 | <0.01 | ** |
|  | *Faecalibacterium* | 1.05 | 0.94 | 0.9 | 0.2 | 0.32 | 0.22 | 0.2 | 0.02 | ** |
|  | *Ruminococcus* | 4.47 | 3.86 | 3.52 | 0.02 | 2.32 | 1.31 | 1.08 | <0.01 | NS |
|  | **Bacteroidetes** | 30.61 | 37.63 | 33.91 | 0.06 | 61.16 | 41.06 | 32.97 | <0.01 | NS |
|  | *Bacteroides* | 16.32 | 18.86 | 17.91 | 0.19 | 17.81 | 12.1 | 8.8 | <0.01 | NS |
|  | *Prevotella* | 4.03 | 4.93 | 4.36 | 0.11 | 32.94 | 21.7 | 16.76 | <0.01 | * |
|  | **Proteobacteria** | 5.57 | 6.32 | 10.84 | 0.01 | 6.97 | 37.53 | 48.63 | <0.01 | NS |
|  | *Acinetobacter* | 0.05 | 0.05 | 2.19 | 0.1 | 0.33 | 0.36 | 18.78 | <0.01 | NS |
|  | *Pseudomonas* | 0.13 | 0.43 | 0.46 | 0.18 | 0.32 | 29.02 | 6.27 | 0.01 | NS |
|  | **Actinobacteria** | 2.69 | 2.46 | 2.43 | 0.01 | 1.82 | 1.29 | 1.31 | 0.01 | ** |
|  | *Bifidobacterium* | 0.35 | 0.31 | 0.29 | 0.13 | 0.27 | 0.16 | 0.14 | <0.01 | NS |
|  | *Slackia* | 0.46 | 0.41 | 0.39 | 0.17 | 0.29 | 0.18 | 0.16 | <0.01 | NS |
|  | **Spirochaetes** | 1.13 | 0.86 | 0.72 | 0.08 | 0.75 | 0.59 | 0.45 | 0.04 | NS |
|  | *Treponema* | 0.62 | 0.37 | 0.31 | 0.11 | 0.35 | 0.29 | 0.22 | 0.11 | NS |
|  | **Fusobacteria** | 0.91 | 0.84 | 0.72 | <0.01 | 0.48 | 0.32 | 0.25 | <0.01 | * |
|  | **Lentisphaerae** | 0.48 | 0.71 | 0.27 | 0.07 | 0.25 | 0.2 | 0.14 | 0.03 | NS |
|  | *Victivallis* | 0.44 | 0.59 | 0.24 | 0.07 | 0.22 | 0.18 | 0.13 | 0.03 | NS |
|  | **Cyanobacteria** | 0.47 | 0.48 | 0.4 | 0.01 | 0.48 | 0.36 | 0.29 | 0.01 | NS |
|  | **Fibrobacteres** | 0.3 | 0.28 | 0.2 | 0.01 | 1.63 | 0.59 | 0.36 | <0.01 | NS |
|  | **Other Bacteria** | 3.07 | 3.18 | 2.56 | 0.07 | 2.74 | 1.95 | 1.48 | <0.01 | NS |
| **Eukaryota** | **unclassified (derived from Eukaryota)** | 0.21 | 0.21 | 0.14 | <0.01 | 0.26 | 0.12 | 0.07 | <0.01 | NS |
|  | **Ascomycota** | 0.12 | 0.1 | 0.09 | <0.01 | 0.12 | 0.08 | 0.06 | 0.01 | NS |
|  | **Streptophyta** | 0.08 | 0.07 | 0.06 | 0.04 | 0.07 | 0.05 | 0.04 | 0.04 | NS |
|  | **Chordata** | 0.05 | 0.06 | 0.05 | 0.24 | 0.07 | 0.08 | 0.09 | 0.16 | * |
|  | **Arthropoda** | 0.01 | 0.01 | 0.02 | 0.46 | 0.04 | 0.03 | 0.02 | 0.02 | NS |
|  | **Other Eukaryota** | 0.15 | 0.14 | 0.13 | 0.01 | 0.16 | 0.11 | 0.11 | 0.05 | NS |
| **Viruses** | **unclassified (derived from Viruses)** | 0.2 | 0.13 | 0.16 | 0.04 | 0.16 | 0.62 | 0.56 | 0.01 | NS |

NS = Not Significant, \* = Significant at 0.05, \*\* = Significant at 0.01
<sup>1</sup>Means are based on 6 cows.

**Fig 2.**
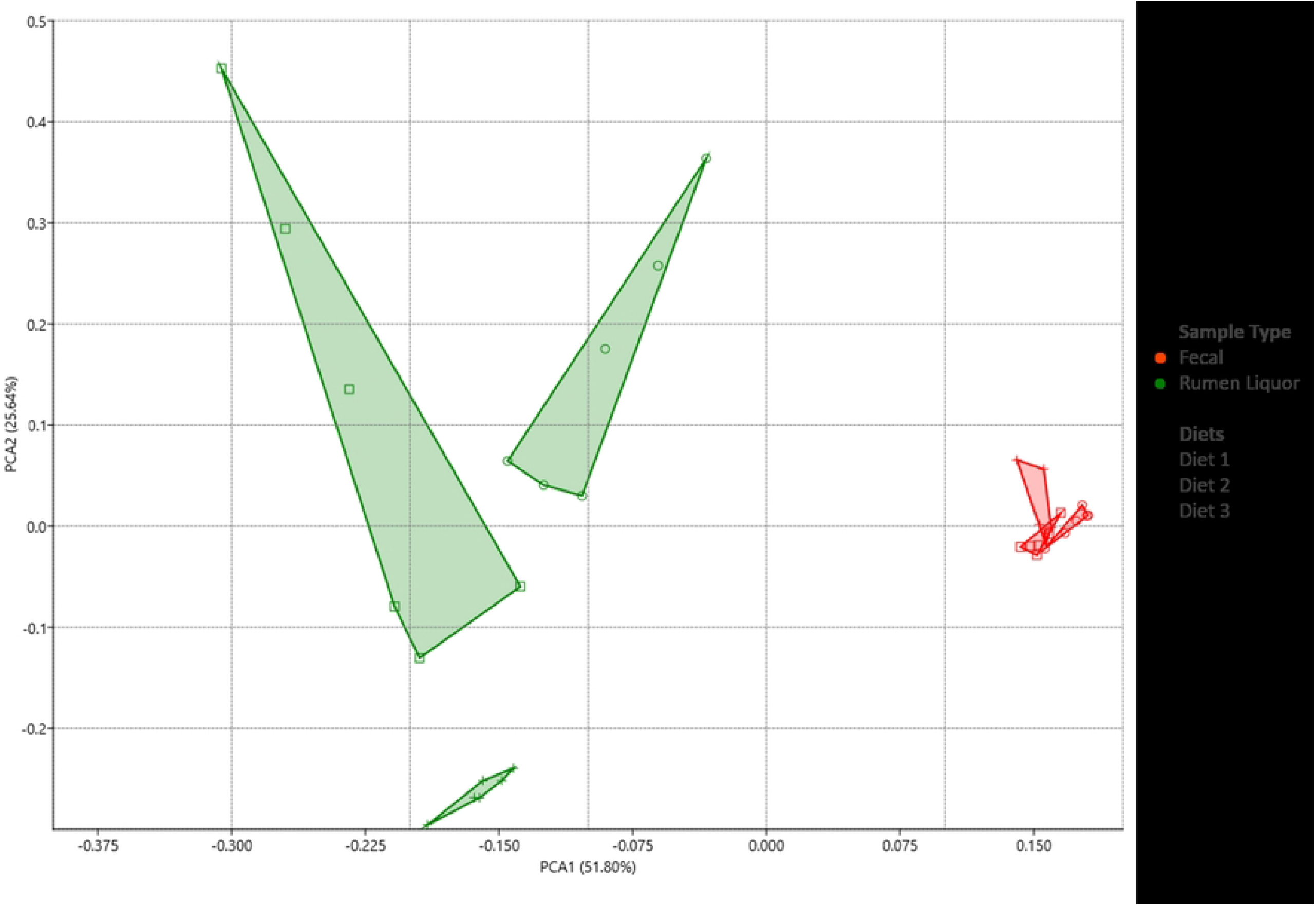
Principal component analysis (PCA) of fecal and rumen liquor microbial communities. The axis of principal component 1 (PC1) described 51.8 % and principal component 2 (PC2) described 25.64 % of total data variability. PCA was performed using PAST v3.13, (35).

Within rumen liquor samples, the ANOVA indicated that all phyla and genera, except the phylum Chordata, significantly differed (p = 0.16) with changes in diet. Diet 1 was associated with higher proportions of all the phyla in the Archaea domain, Firmicutes (p = 0.01), Bacteroidetes (p < 0.01), Actinobacteria (p = 0.01), Spirochaetes (p = 0.04), Fusobacteria (p < 0.01), Lentisphaerae (p = 0.03), Cyanobacteria (p = 0.01), Fibrobacteres (p < 0.01), other minor bacterial phyla (p = 0.01), unclassified derivatives from Eukaryota (p < 0.01), Ascomycota (p = 0.01), Streptophyta (p = 0.04), and other minor Eukaryotic phyla (p = 0.05), whereas Diet 3 had the lowest proportions in the same phyla. Conversely, Proteobacteria (p < 0.01) and unclassified derivatives of viruses (p = 0.01) had the highest and lowest abundance in Diet 3 and Diet 1, respectively. All the featured genera showed a similar abundance distribution at the genus level as that of their phyla with different diets (Table 2). The shift observed in the three most abundant phyla was important as the animals were moved from a high-fiber diet (Diet 1) to a high-concentrate diet (Diet 3). Firmicutes and Bacteroidetes showed a consistent decrease in abundance, whereas Proteobacteria showed a consistent increase from Diet 1 to Diet 3 (Table 2). The most abundant bacteria species were *Prevotella ruminicola*, and it showed a significant decrease with an increased level of concentrate in the diet. The other bacterial species identified in this study did not show significant variations with changes in diet.

### Variation in microbial taxa in fecal and rumen liquor sample types

Given that obtaining rumen liquor samples is more tedious and invasive than collecting fecal samples, we investigated the theory that the fecal metagenome of a cow could be used as a predictor of its rumen metagenome. A comparison of alpha diversity indices revealed that rumen microbiota had a higher species richness than fecal microbiota, as indicated by the Chao 1 index. Conversely, fecal microbiota was more diverse and had greater evenness than rumen microbiota (P < 0.05; Table 1). Similarly, fecal and rumen liquor samples were separated into two clusters based on PCA analysis (Fig 2). For further analysis, we compared the relative abundance of the most common phyla (relative abundance > 0.1 %) and genera (relative abundance > 0.05 %) between fecal and rumen liquor samples (Table 2). The relative abundance values were the mean abundance of all the treatments and animals within each sample type. Relative abundance revealed that four phyla, Firmicutes (p < 0.01), Actinobacteria (p < 0.01), Euryarchaeota (p = 0.03), and Fusobacteria (p = 0.01), had significant differences between fecal and rumen liquor samples. There were significant differences between fecal and rumen liquor sample types in five genera (Table 2, Fig 3).

**Fig 3.**
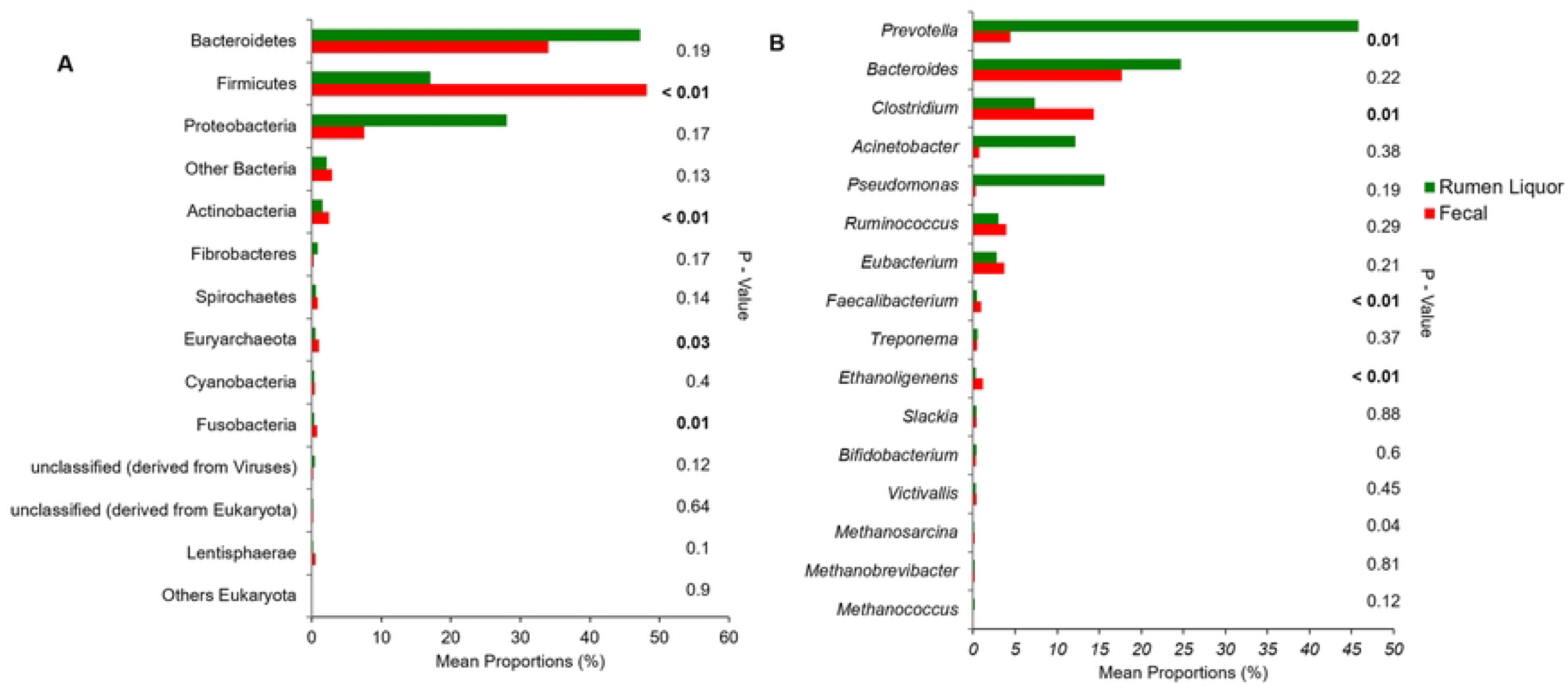
Comparison of relative abundance of microbial phyla and genera in fecal and rumen liquor sample types. The 15 most abundant phyla with relative abundance > 0.1 in both fecal and rumen liquor sample types. B The 17 most abundant genera with relative abundance > 0.05 in both fecal and rumen liquor samples.

Moreover, PCA revealed the clustering of fecal samples away from rumen liquor samples, particularly in the PC1 dimension. The first two components of the Bray-Curtis dissimilarity matrix’s non-metric multidimensional scaling showed a significant distance difference between the two sample types (p < 0.01, from 9999 permutations of the one-way PERMANOVA) (Fig 2). Furthermore, a comparison of the entire OTU dataset by linear discriminant analysis effect size (LEfSe) identified 847 features to be significantly discriminative between fecal and rumen liquor samples. Of these significantly discriminative features, 30 had an absolute LDA score > 4.0. One phylum, Firmicutes, was differentially abundant for fecal samples, while Bacteroidetes and Proteobacteria were much more enriched in rumen liquor samples (Fig 4A). The genera, *Prevotella* and *Clostridium*, were the highest sources of variation between the communities, with an absolute LDA score factor of roughly 4.8 (Fig 4B). To test if the fecal microbial profile of a cow could predict the rumen liquor metagenome profile, the correlation between each fecal and rumen liquor metagenome profile was determined. These correlations were then evaluated to determine if they were greater for samples from the same animal than between animals. We observed no significant differences between correlations from the same animal and those from between animal samples (t-test; p = 0.914) (Table 3).

**Fig 4.**
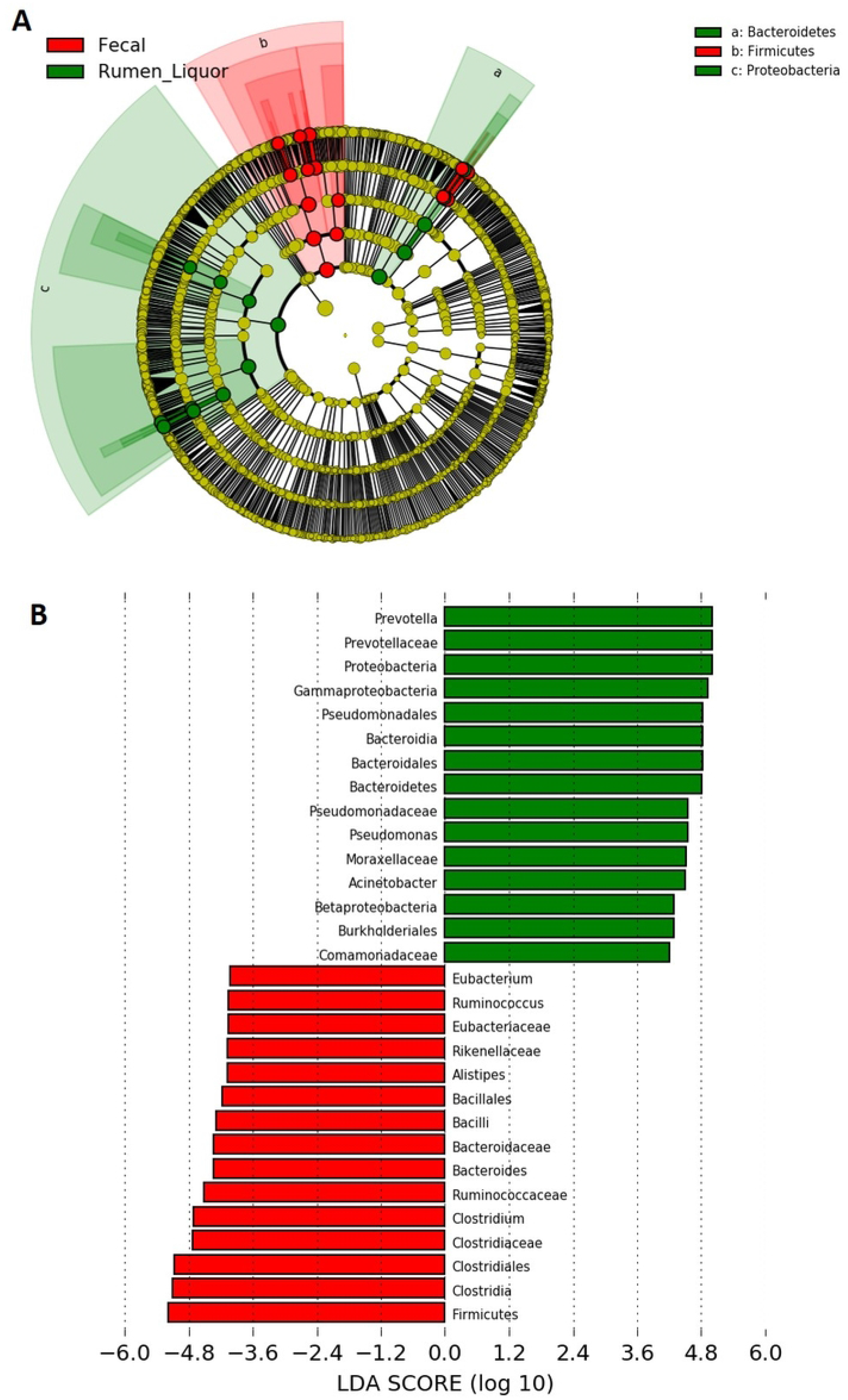
LEfSe analysis. (**A**) The cladogram illustrating the presence of bacterial phyla that are significantly different between the fecal and rumen liquor samples types. (B) The histogram of the Linear discriminant analysis (LDA) scores illustrating the differentially abundant microbial communities in the rumen (green) and fecal (red) sample types. The LDA score at log 10 > 4 is set as threshold and the length of each bin, i.e., LDA score represents the extent to which the microbial taxa differ among the groups.

**Table 3:**
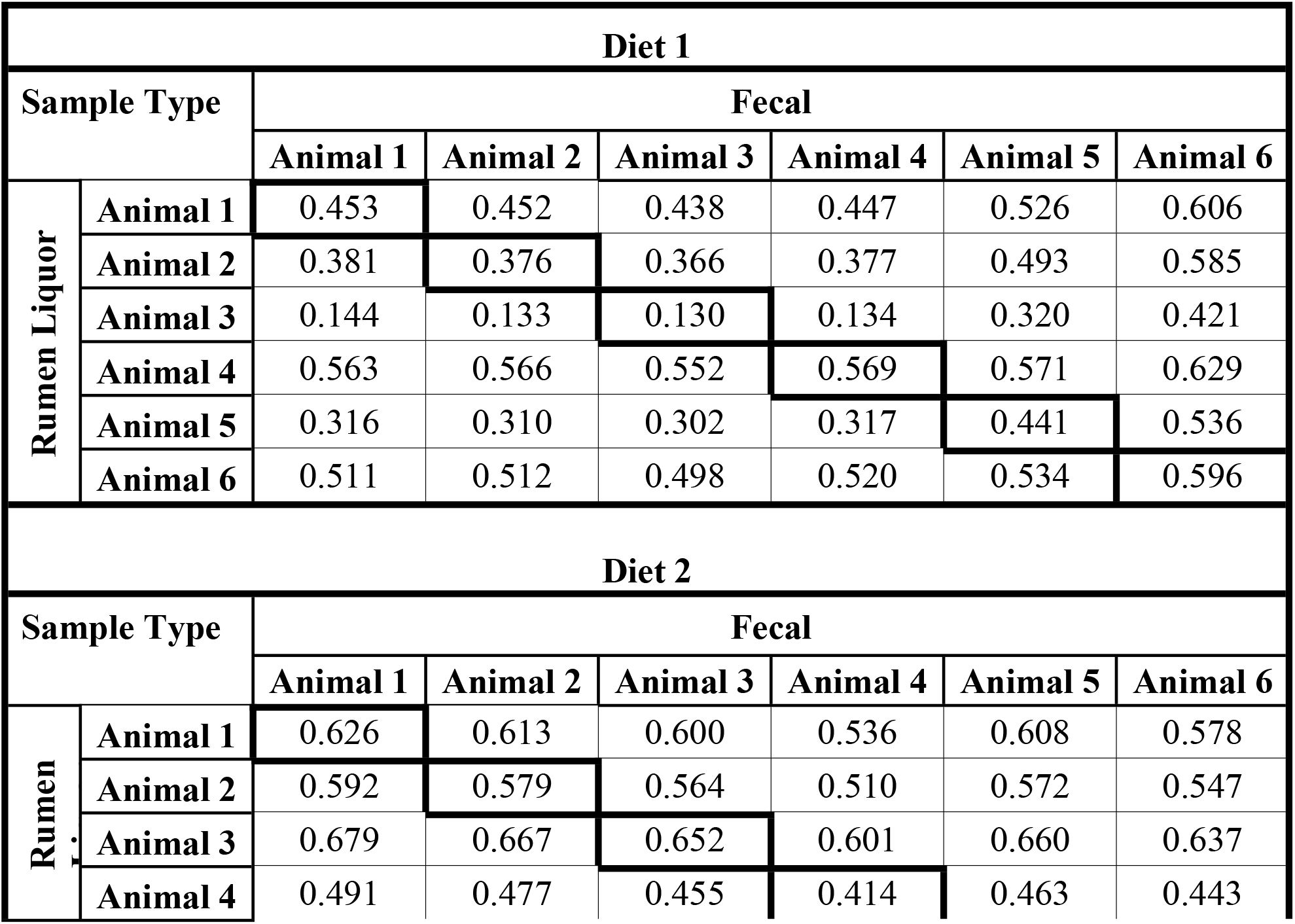
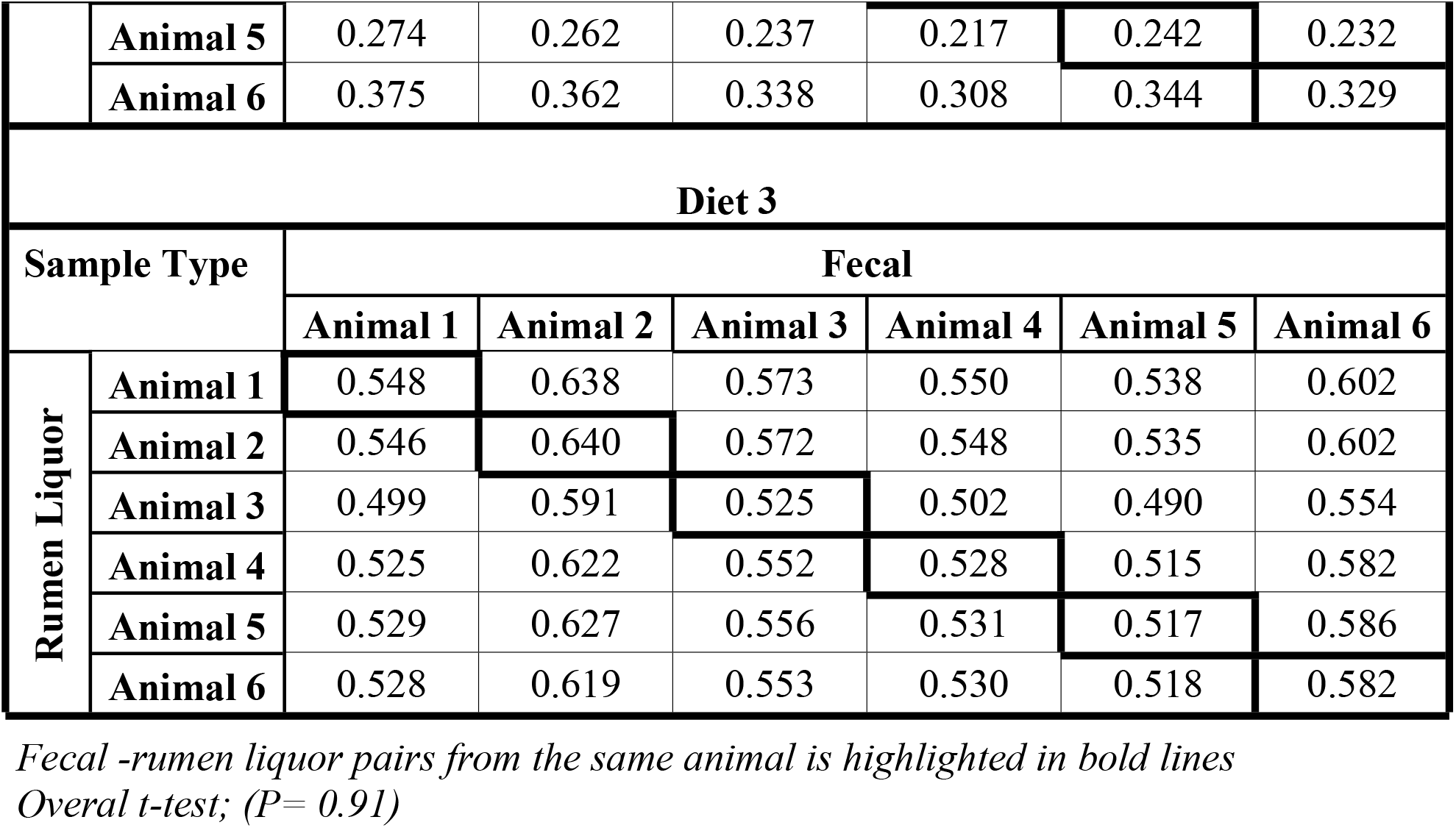
Correlation between fecal-rumen liquor pairs.

### Functional Capacity of Ruminal and Fecal Microbiota

The analysis of enzyme functions gives the microbial community a metabolic blueprint. We used the SEED subsystems classification in the MG-RAST pipeline to assess if the sequence fragments were associated with specific metabolic functions. Normalization was carried out before the metabolic potential among the samples was compared. Normalization was done to account for differences in community structure, library size and to compare functional categories with low abundance (42). The normalization was performed together for all pathways at Level 1 and within each specific pathway at level 2 classification. Table 4 illustrates the 28 most abundant predicted functional pathways found at level 1 classification. The most abundant level 1 pathway was carbohydrate metabolism ranging from 14.08 % to 14.19 % and 12 % to 14.62 % in fecal and rumen liquor samples, respectively, followed by clustering-based subsystems (functional coupling evidence but unknown function; fecal: 13.97 % to 14.28 %, rumen liquor: 11.99 % to 13.21 %), and then protein metabolism (12.5% to 13.37% in fecal and 8.67 % to 11.67% in rumen liquor samples). Other dominant pathways were amino acids and derivatives (8.78–10.13 %) and miscellaneous (5.52–6.14 %), suggesting the dominant role of these functional pathways in all the samples. ANOVA revealed that dietary changes caused significant variation in only two pathways changes within fecal samples, whereas changes in diet resulted in significant variations in all the functional pathways, except miscellaneous (p = 0.152), in rumen liquor samples. All pathways were significantly varied between fecal and rumen sample types except DNA metabolism (p = 0.057) and photosynthesis (p = 0.279) (Table 4).

**Table 4:** Relative abundance (%) of Level 1 subsystems classification of microbial functional pathways.

| Functional Pathway | Fecal<sup>1</sup> |  |  |  | Rumen Liquor<sup>1</sup> |  |  |  | Sample Type<br>P value |
| --- | --- | --- | --- | --- | --- | --- | --- | --- | --- |
|  | Diet 1 | Diet 2 | Diet 3 | P value | Diet 1 | Diet 2 | Diet 3 | P value |  |
| Amino Acids and Derivatives | 9.018 | 8.89 | 8.776 | 0.135 | 9.233 | 10.13 | 9.956 | 0.001 | *** |
| Carbohydrates | 14.111 | 14.191 | 14.079 | 0.878 | 14.621 | 12.27 | 12 | <0.001 | * |
| Cell Division and Cell Cycle | 1.328 | 1.444 | 1.539 | 0.008 | 1.101 | 0.924 | 0.997 | 0.013 | *** |
| Cell Wall and Capsule | 4.245 | 4.329 | 4.287 | 0.779 | 5.846 | 5.555 | 5.17 | 0.001 | *** |
| Clustering-based subsystems | 13.971 | 14.191 | 14.286 | 0.395 | 13.211 | 12.527 | 11.988 | <0.001 | *** |
| Cofactors, Vitamins, Prosthetic Groups, Pigments | 3.874 | 3.759 | 3.571 | 0.008 | 4.274 | 4.499 | 4.868 | <0.001 | *** |
| DNA Metabolism | 6.1 | 6.775 | 6.653 | 0.002 | 6.973 | 5.81 | 5.327 | <0.001 | NS |
| Dormancy and Sporulation | 0.764 | 0.833 | 0.926 | 0.059 | 0.309 | 0.245 | 0.197 | 0.004 | *** |
| Fatty Acids, Lipids, and Isoprenoids | 1.636 | 1.592 | 1.567 | 0.615 | 1.596 | 2.081 | 2.3 | <0.001 | *** |
| Iron acquisition and metabolism | 0.609 | 0.43 | 0.429 | 0.001 | 0.683 | 2.093 | 1.491 | <0.001 | *** |
| Membrane Transport | 3.133 | 2.623 | 2.727 | <0.001 | 2.571 | 3.876 | 3.879 | <0.001 | ** |
| Metabolism of Aromatic Compounds | 0.458 | 0.289 | 0.261 | 0.002 | 0.353 | 0.892 | 1.332 | <0.001 | *** |
| Miscellaneous | 5.52 | 5.61 | 5.531 | 0.616 | 5.92 | 6.138 | 6.052 | 0.152 | *** |
| Motility and Chemotaxis | 0.388 | 0.24 | 0.313 | 0.021 | 0.387 | 1.017 | 1.221 | <0.001 | *** |
| Nitrogen Metabolism | 0.43 | 0.29 | 0.276 | 0.001 | 0.839 | 0.859 | 1.125 | <0.001 | *** |
| Nucleosides and Nucleotides | 4.607 | 4.526 | 4.393 | 0.225 | 3.905 | 3.492 | 3.613 | 0.007 | *** |
| Phages, Prophages, Transposable elements, Plasmids | 2.157 | 1.919 | 2.28 | 0.069 | 2.437 | 3.004 | 2.932 | 0.271 | *** |
| Phosphorus Metabolism | 0.668 | 0.541 | 0.603 | 0.006 | 0.601 | 0.77 | 0.965 | <0.001 | *** |
| Photosynthesis | 0.02 | 0.019 | 0.016 | 0.792 | 0.014 | 0.011 | 0.02 | 0.086 | NS |
| Potassium metabolism | 0.288 | 0.213 | 0.24 | 0.072 | 0.339 | 0.53 | 0.572 | <0.001 | *** |
| Protein Metabolism | 12.497 | 13.425 | 13.365 | 0.109 | 11.671 | 9.114 | 8.662 | <0.001 | *** |
| RNA Metabolism | 5.387 | 5.838 | 5.893 | 0.003 | 5.121 | 4.481 | 4.411 | <0.001 | *** |
| Regulation and Cell signaling | 0.827 | 0.645 | 0.655 | <0.001 | 0.747 | 1.136 | 1.184 | 0.001 | *** |
| Respiration | 3.279 | 3.151 | 3.07 | 0.219 | 2.662 | 2.671 | 3.268 | <0.001 | ** |
| Secondary Metabolism | 0.085 | 0.04 | 0.044 | 0.015 | 0.114 | 0.121 | 0.201 | <0.001 | **** |
| Stress Response | 1.814 | 1.643 | 1.667 | 0.031 | 1.616 | 2.085 | 2.368 | <0.001 | ** |
| Sulfur Metabolism | 0.569 | 0.439 | 0.462 | 0.017 | 0.595 | 0.971 | 1.006 | <0.001 | *** |
| Virulence, Disease and Defense | 2.216 | 2.115 | 2.092 | 0.248 | 2.265 | 2.698 | 2.896 | 0.005 | *** |

NS = Not Significant, \* = Significant at 0.05, \*\* = Significant at 0.01, \*\*\* = Significant at 0.001
<sup>1</sup>Means are based on 6 cows.

A total of 168 level 2 pathways were identified in the rumen and feces samples. Further detailed, level 2 analysis was undertaken on carbohydrates and protein metabolism pathways as they were the most abundant functional pathways at level 1 classification. The carbohydrate metabolism included eleven level 2 pathways. Amino sugars, central carbohydrate metabolism, fermentation, one-carbon metabolism, and organic acids significantly changed in fecal samples when the diet was varied. Conversely, only one pathway, CO_2_ fixation (p = 0.295), did not vary across diets in rumen liquor samples. Central carbohydrate metabolism, fermentation, organic acids, and sugar alcohols pathways increased with an increase in concentrate, whereas monosaccharides, disaccharides, oligosaccharides, and polysaccharides decreased with a greater proportion of dietary concentrate (S2 Table). Five different pathways, protein biosynthesis, protein degradation, protein folding, protein processing and modification, and selenoproteins were identified at level 2 classification of protein metabolism pathway. The maximum sequences were attributed to protein biosynthesis, followed by protein degradation. Protein degradation (p = 0.01) and selenoproteins (p = 0.001) were the only categories that varied in fecal samples. The variation in protein degradation did not follow any specific dietary pattern, whereas selenoproteins increased with increased concentration in the diet. Within the rumen liquor samples, the abundance of all the categories varied across the diets.

With an increase in concentrate level, all the categories except protein biosynthesis increased (S2 Table). MG-RAST-BLAT integration revealed that carbohydrate metabolism pathways were spread across 41, 333 and 28 genera from archaea, bacteria, and Eukaryota domains, respectively. Protein metabolism pathways were generally scattered across 43 archaea, 341 bacteria, and 38 Eukaryota microbial genera. An assessment of the microbes responsible for the most abundant level 2 pathways (protein biosynthesis and central carbohydrate metabolism) revealed that within these two pathways, and irrespective of the diet, the prominent genera were *Bacteroides* and *Clostridium* in fecal samples. However, in rumen liquor samples, Diet 1 was dominated by *Bacteroides* and *Clostridium*; Diet 2 was dominated by *Bacteroides* and *Pseudomonas;* and Diet 3 was dominated by *Acinetobacter, Bacteroides*, and *Pseudomonas* (Table 5).

**Table 5:**
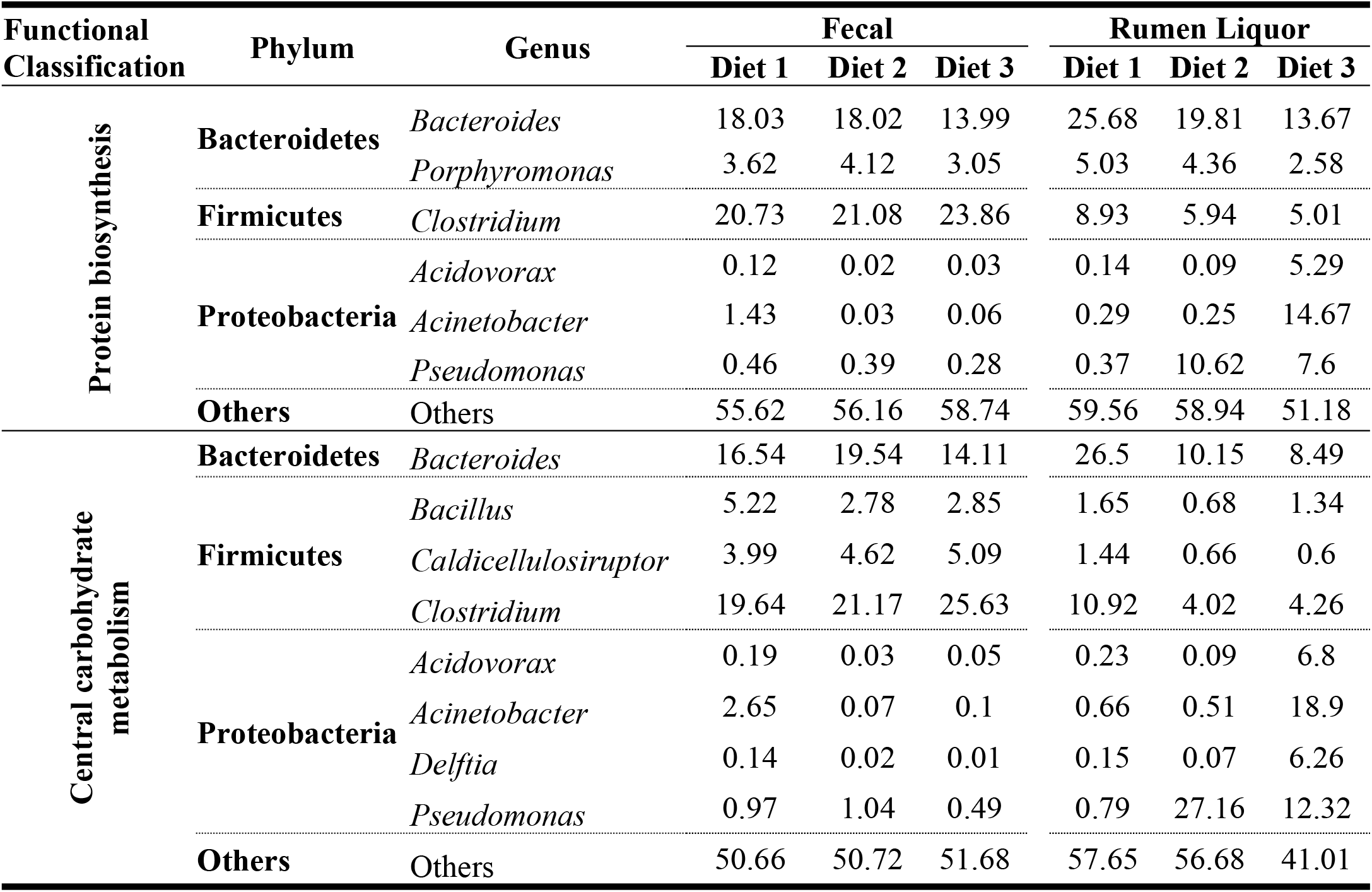
Relative abundance (%) of predominant microbiota associated with protein biosynthesis and central carbohydrate metabolism.

Despite their low normalized abundance at level 1 classification, a detailed assessment was carried out on sulfur metabolism, stress response, cell wall and capsule, and dormancy and sporulation functional pathways. This assessment was because previous studies have shown the importance of sulphur in rumen microbial synthesis. Moreover, the rumen microbiome goes through an array of dietary stresses. To adapt to these constraints, rumen microbes have developed several stress responses like the capacity to enter into dormant states (spores) or a cell wall/capsule development. For the fecal samples, only stress response (p = 0.031) and sulfur metabolism (p = 0.017) were significantly affected by dietary changes; however, all the pathways were significantly influenced by dietary variations for rumen liquor samples. Changes in pathways did not follow any variation in the diet for fecal samples, while in the rumen liquor samples, all the pathways except cell wall and capsule decreased with an increase in concentrate proportion in the diet.

In the sulfur metabolism pathway, the relative abundance of the level 2 functional category of inorganic sulfur assimilation reduced with increased concentration in the diet. However, organic sulfur assimilation increased with increased concentrations in the diet (S2 Table). The OTUs involved in sulfur metabolism belonged to several genera, as shown in Table 6. The stress response pathway contained seven categories at level 2 classification. The most abundant pathways in the rumen and feces were oxidative stress, osmotic stress, and heat shock, whereas acid stress, periplasmic stress, cold shock, and desiccation stress had the least abundance. The diet had significant effects in all the pathways within fecal samples except desiccation stress (p = 0.035) and periplasmic stress (p = 0.063). Acid stress (p = 0.798) and desiccation stress (p = 0.155) were the only two pathways that were not significantly affected by diet in rumen liquor samples.

**Table 6:**
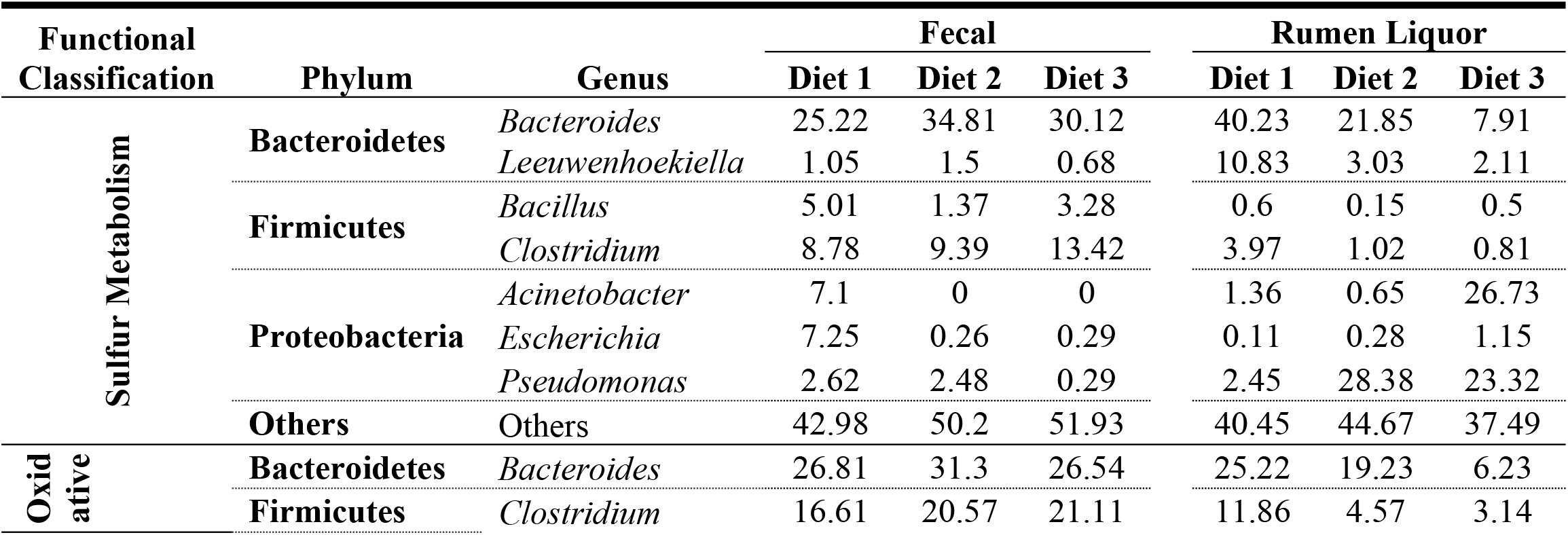
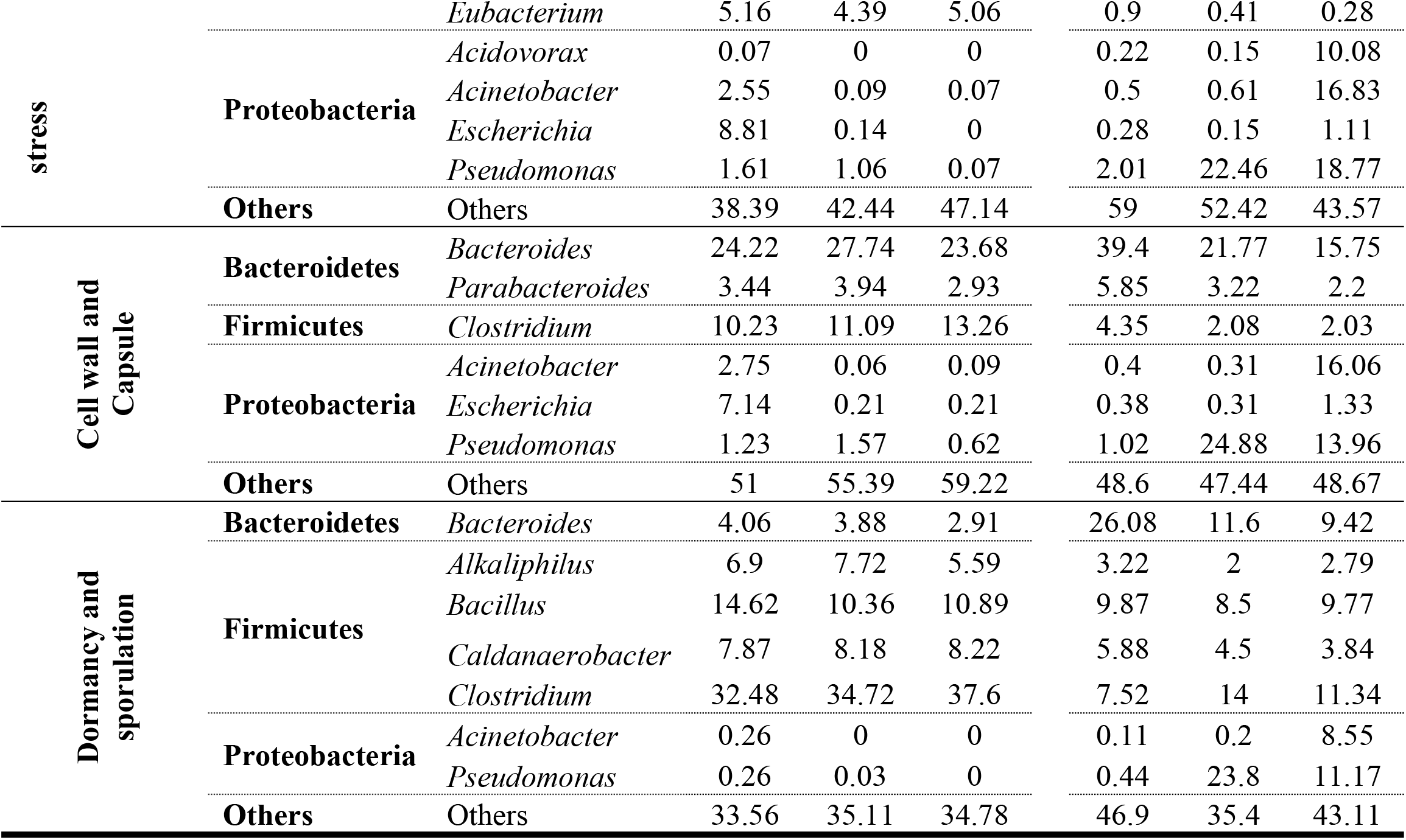
Relative abundance (%) of predominant microbiota associated with sulfur metabolism, oxidative stress, cell wall and capsule and dormancy and sporulation pathways.

The pathways that were significantly affected by dietary changes did not follow any dietary trend in fecal samples. Conversely, in rumen liquor samples, osmotic stress, oxidative stress, and cold shock increased with increased concentration. This was in contrast to heat shock, which decreased with an increase in concentrate proportion (S2 Table). A total of 31 phyla and 372 genera spread across archaea, bacteria, and Eukaryota domains were affiliated with the stress response pathways. The results further showed that for oxidative stress response, *Bacteroides* and *Clostridium* were the most prominent genera, irrespective of the diet, in the fecal samples. In contrast, *Bacteroides* dominated Diet 1, Bacteroides and Pseudomonas were the highest genera with Diet 2, and genera Acidovorax, Acinetobacter, and Pseudomonas were the highest dominant with Diet 3 (Table 6).

Four pathways were identified in level 2 classification of the cell wall and capsule in all the samples. These pathways were capsular and extracellular polysaccharides, the cell wall of mycobacteria, gram-negative cell wall components, and gram-positive cell wall components. The gram-negative cell wall components increased with an increasing amount of dietary concentrate, whereas the capsular and extracellular polysaccharides reduced. Genus-wise affiliation of the most abundant genera associated with the cell wall and capsule pathways were similar to those observed for the sulfur metabolism category, as shown in Table 6. Level 2 pathways were associated with spore DNA protection and unknown (null) function within the dormancy and sporulation pathway. Changes in diet did not exert significant changes in these pathways for both fecal and rumen liquor samples (S2 Table). *Clostridium* was the dominant genus involved in dormancy and sporulation for all fecal samples, while several diverse genera were associated with different diets in rumen liquor samples (Table 6).

## Discussion

Inadequate feed resources are one of the greatest challenges facing dairy farmers in the East African tropics. This problem is aggravated by the high cost of commercially formulated feeds. For these reasons, dairy farmers subject their animals to constant changes in the quality and quantity of feeds (43). These feed resources, which consist mainly of indigestible plant cell wall components, are degraded by rumen microbes into volatile fatty acids and microbial proteins that are critical for the hosts’ survival and production (44). Among other factors, diet is a major driver of shifts in cattle GI tract microbial communities (10,45,46); however, information on the shifts in rumen GI tract microbiota of crossbred cattle reared by dairy farmers in the East African tropics is limited. Additionally, given the difficulty in collecting rumen liquor samples from cattle, it would be expedient if there was a relationship between the microbial structure of rumen fluid and other samples that are easy to obtain, like feces. In this study, we characterized shifts in ruminal and fecal microbiota and associated microbial functional roles occurring due to increasing dietary concentrate proportion. Additionally, we assessed the suitability of using fecal metagenome as a proxy for rumen metagenome in crossbred dairy cattle.

### Microbial diversity and taxonomic assessment

Our results indicated that dietary changes influenced the community composition and abundance of several rumen microbial taxa, while very few OTUs were varied in the feces. Similar to previous studies, (47–50), in rumen liquor samples, the microbial richness decreased with an increase in concentrate proportion in the diet. This observation could have been due to extensive fermentation activities and a low rumen pH due to increased easily fermentable nutrients from the higher dietary concentrate proportion (51). Thus this environment was perhaps less favorable for some members of rumen microbiota, and as such, microbial richness declined.. PCA-based clustering analysis showed differences in treatments within rumen liquor samples (Fig 4). Moreover, PCA results showed that Diet 1 separated significantly from Diet 2 and Diet 3. We hypothesize that this observation may have resulted from the relatively higher abundance of OTUs, especially from Bacteroidetes phylum, recovered when the animals were fed on a diet with high roughage proportion (Diet 1).

As reported in previous studies of cattle (37–39), the most abundant phyla were Bacteroidetes, Firmicutes, and Proteobacteria, irrespective of sample type and diet. Similar to findings reportrd by Fernando *et al*. (47), changes from a high-roughage diet (Diet 1) to a high-concentrate diet (Diet 3) caused a consistent rise in the abundance of Proteobacteria and a decrease in the abundance of Bacteroidetes and Firmicutes in rumen liquor samples. Previous studies have demonstrated that members of the Bacteroidetes phylum can utilize starch, xylan, pectin, galactomannan, and arabinogalactan (52) because they have higher mean polysaccharide lyases (PLs) and glycoside hydrolases (GHs) genes per genome, as well as signal peptide-containing PLs and GHs, compared to members of any other bacterial phyla in the GI tract (53). As such, members of the Bacteroidetes phylum are among the primary degraders of the many complex polysaccharides in the plant cell wall (54). The Firmicutes, on the other hand, can utilize carbohydrates such as xylan, cellulose, hemicellulose, and galactomannan as energy sources (Dassa et al., 2014; Morrison and Miron, 2000). This explained their higher abundance when animals were fed on Diet 1 and subsequently decreased as the concentration increased. Conversely, increased Proteobacteria abundance with an increase in concentrate proportion in the diet suggests a greater need for bacteria to digest the newly available fermentable carbohydrates (47,49). Moreover, when animals are fed higher amounts of starchy feed, the competitiveness of the members of Bacteroidetes phyla in the microbial community declines, allowing opportunistic phyla, such as Proteobacteria, to proliferate faster per unit of time, resulting in an increase in the proportions of Proteobacteria (57,58).

The most abundant genera were *Prevotella, Bacteroides*, and *Clostridium*. While the abundance of these genera was relatively stable in fecal samples, their abundance decreased with an increase in concentrate proportion in rumen liquor samples. The predominance of these genera can be explained in part because i) *Prevotella* genus is comprised of genetically and metabolically diverse members (59) that are numerically high in animals fed on both high-grain and high-forage diets (60,61). *Prevotella* species can use starches, simple sugars, and other non-cellulosic polysaccharides as energy sources (62). Furthermore, *Prevotella* species, including members with hemicellulolytic and proteolytic activity (61), are considered to be involved in hemicellulose and pectin digestion (63) and protein or peptide metabolism (64,65) in the rumen. ii) The genus *Bacteroides* has been shown to extensively contribute to carbohydrate, small molecule, and organic acid metabolisms and plays a significant role in other Bacteria-linked metabolic processes (66). iii) *Clostridia* contains cellulolytic strains that are mainly commensals in the GI tract. *Clostridia* members make up a substantial part (10-40%) of the total bacteria in the gut microbiota (67,68). As such, Clostridia likely plays a crucial role in gut homeostasis by interacting with the other resident microbe populations and providing specific and essential functions (69).

Dietary variations significantly affected very few OTUs in fecal samples. This was further supported by a distinct clustering trend in all fecal microbiota samples. This was in congruence with a previous study that characterized rumen and fecal microbiome in bloated and non-bloated cattle (70). The few OTUs that were significantly affected by diet did not follow any pattern corresponding with the diets as previously shown by (10,45,47,71). The stability of the fecal metagenome with changes in diet observed in this study could be attributed to the stability of the hindgut, especially in its pH, thus promoting greater substrate availability and consequently microbial stability (72).

It is challenging to obtain rumen liquor samples; hence, it would be advantageous if the microbe profiles of the rumen and other samples, such as fecal samples, which can be easily collected, are considerably overlapped. Given the depth and precision of NGS technology in assessment of microbial populations, we hypothesized that any relationship, however small, between rumen and fecal microbiota would be identified. We could not find proof of a significant link between rumen liquor and fecal profiles, similar to earlier studies that assessed the metagenome profiles of rumen liquor and fecal samples (11,24,73). The dissimilarity in fecal and rumen liquor samples was shown by fecal microbiota being more diverse and having greater evenness than rumen microbiota. Moreover, there was sample type separation observed from the results of PCA analysis and a significant difference in the abundance of a large number of OTUs of fecal and rumen liquor samples. Similar to past studies that showed a significant difference in cattle microbiota in various sections of the GI tract (74,75), LEfSe analysis showed different OTUs being differentially selected in fecal and rumen liquor samples. The differences between fecal and rumen liquor metagenomes may be largely related to the role of the two environments; the rumen microbiota may be intensely selected to remain functional as the host relies on them for digestion, while the fecal metagenome may be less restricted (11). Additionally, luminal pH, differences in gut motility, host secretions, and nutrients are other factors that affect the microbial structure in various sections of the gut (76). We also found no indication to support our hypothesis that the fecal microbial profile of a cow correlated more closely with that of its rumen liquor than that of any other cow. Although this observation was in agreement with an earlier study (11), more advanced analysis and additional sample types may be needed to explore this theory.

### Functional metagenomic classification

Shotgun sequencing provides the ability to classify microbes taxonomically and allows for assessing microbial functional pathway analysis by annotation to remote databases (77). In some ways, this distribution of functional pathways enables the identification of the characteristics of substrate utilization by the rumen ecosystem (78). In the present study, fecal and rumen liquor sample types showed a significant variation in functional classification between the various diets. The largest percentage of gene functions were linked to protein and carbohydrate metabolism which are vital for microbial survival and proliferation (79,80). The high abundance of carbohydrate and protein metabolism functional pathways was consistent with the findings of previous studies on cattle (75), humans (81), and mice (82). Level 2 classification of protein metabolism pathway revealed that most ontologies were associated with protein biosynthesis. This high abundance of protein biosynthesis pathway could be explained in part by 19 different tRNA aminoacylation categories for different amino acids identified on further (level 3) analysis of this category. When the effect of diet was tested on level 2 pathways in protein metabolism, differences in metabolic potential were observed, indicating a selective pressure exerted on the microbes with particular metabolic capabilities. For example, the selenoproteins metabolism pathway increased with an increased amount of concentrate in the diet. Selenoproteins are involved in combating oxidative stress (83). As seen in this study, an increase in concentrate concentration caused an increase in oxidative stress (84). Consequently, selenoproteins pathway increased to counter this increase in oxidative stress. In carbohydrate metabolism, the increasing levels of central carbohydrate metabolism and fermentation pathways with the expected higher content of easily available carbohydrates in a high-concentrate, low-roughage diet.

Our results reported an increase in sulphur metabolism as the animals were transitioned from a high-fiber to a high-concentrate diet. We theorize that the high amount of concentrate led to a reduction of members of rumen microbiota due to a less favorable environment as a result of lactate accumulation. To cater for this, sulphur metabolism was increased to improve microbial growth and consequently utilization of lactate by the rumen bacteria by contributing to the synthesis of amino acids, especially methionine and cysteine (85). This theory was supported by an apparent increase, although not significant, of the level 2 Lysine, threonine, methionine, and cysteine pathway within the Amino Acids and Derivatives metabolism pathway.

Microbes survive in the rumen under different stresses, which may be natural or associated with the feed. Some of the feed-associated stresses, such as anti-nutritional factors, act as natural antimicrobial agents by limiting the growth of some microbes (34). To adapt to these constraints, rumen microbes, especially bacteria, can enter into dormant states (spores) or develop a cell wall/capsule (86). Spores serve to protect the bacterium from harmful environmental conditions by reducing it into a desiccated, cryptobiotic and highly defensive state which conveys resistance to many environmental assaults that would otherwise harm and kill the vegetative form of the bacterium (34). We theorize that the increase in the relative abundance of stress response and gram-negative cell wall component pathways with an increase in the amount of concentrate diet was a response to external stress experienced by the gut microbial community caused by dietary changes. Further, the identification of spore DNA protection suggests the potential for long-term dormancy of cells through DNA protection.

However, it is crucial to highlight that functional prediction is based on known roles of the microbial communities found in the GI tracts of humans and animals as determined by the whole genome shotgun sequencing of these samples. Due to the scarcity of shotgun sequencing research in ruminants, the functionality of ruminant GI microbiota members may be overestimated or underestimated (87).

## Conclusion

This study compared the diversity and functional roles of fecal and rumen liquor microbial communities in crossbred cows under different diets. Our findings indicate that dietary modifications had a significant effect on several rumen and fecal microbial OTUs. In the rumen, an increase in dietary concentrate amount resulted in an upsurge in the abundance of Proteobacteria, while reducing the abundance of Bacteroidetes and Firmicutes. Conversely, the changes in microbial composition in fecal samples were not consistent with the dietary modification patterns. Thus, there was no significant relationship between the rumen and fecal metagenome. Moreover, fecal microbiota from one animal did not correlate more than that from different animals. Functional classification identified that microbial genes were dominated by those associated with carbohydrate metabolism and protein metabolism. The assessment of dietary effects on microbial functions revealed that an increased roughage level in the diet boosted protein synthesis while decreasing central carbohydrate metabolism. This study identified that *Bacteroides, Clostridium*, and *Pseudomonas* genera were the principal hosts of these microbial functions. Our current study suggests a connection between the feed of the host dairy cattle and their resident rumen microbiota. The study also reveals potential candidate taxa that may prove useful for future inoculation studies given their association with either roughage or concentrate based diets. However, additional work is needed to evaluate other potential sample types that can serve as proxies for rumen microbial composition and functional capacity in crossbred dairy cattle.

## Acknowledgements

The BecA-ILRI Hub supported this work through the Africa Biosciences Challenge Fund (ABCF) program. The ABCF program is funded by the Australian Department for Foreign Affairs and Trade (DFAT) through the BecA CSIRO partnership; the Syngenta Foundation for Sustainable Agriculture (SFSA); the Bill & Melinda Gates Foundation (BMGF); the UK Department for International Development (DFID) and the Swedish International Development Cooperation Agency (Sida).

## Supporting information

**S1 Text: Supporting information, containing tables on diet formulation, chemical analysis and metagenomic sequence sample details**. Table A. Roughage-based diet supplemented with dairy meal concentrate. Table B. Chemical composition of the dietary components. Table C. Analysis of Variance (ANOVA) for the 36 metagenomic sequence sample details

**S2 Table. Normalized mean abundances for selected SEED Level 2 functional categories of diets within fecal and rumen liquor samples**

